# A Novel Taxonomic Database for eukaryotic Mitochondrial Cytochrome Oxidase subunit I Gene (eKOI): Enhancing taxonomic resolution at community-level in metabarcoding analyses

**DOI:** 10.1101/2024.12.05.626972

**Authors:** Rubén González-Miguéns, Alex Gàlvez-Morante, Margarita Skamnelou, Meritxell Antó, Elena Casacuberta, Daniel J. Richter, Daniel Vaulot, Javier del Campo, Iñaki Ruiz-Trillo

**Affiliations:** Institut de Biologia Evolutiva (CSIC-Universitat Pompeu Fabra), 08003, Barcelona, Spain; ICREA, Barcelona, Spain; Sorbonne Université, CNRS, UMR7144, Station Biologique de Roscoff, 29680, Roscoff, France; Department of Biosciences, University of Oslo, PO Box 1066 Blindern, 0316, Oslo, Norway

## Abstract

Metabarcoding has emerged as a robust method for understanding biodiversity patterns by retrieving environmental DNA (eDNA) directly from ecosystems. Its low cost and accessibility have extended its use across biological topics, from symbiosis to biogeography, and ecology. A successful metabarcoding application depends on accurate and comprehensive reference databases for proper taxonomic assignment. The 18S rRNA gene is the primary genetic marker used for general/broad eukaryotic metabarcoding due to its combination of conserved and hypervariable regions, and the availability of extensive taxonomically-informed reference databases like PR2 and SILVA. Despite its advantages, 18S rRNA has certain limitations at lower taxonomic levels, depending on the lineage. Alternative fast-evolving molecular markers, such as the mitochondrial cytochrome oxidase subunit I (COI) gene, have been adopted as widely used "barcoding genes" for eukaryotes due to their resolution to the species level. However, the COI gene lacks a curated taxonomically-informed database covering all eukaryotes, including protists, comparable to those available for 18S rRNA. To address this gap, we introduce eKOI, a curated COI gene database aimed at enhancing the taxonomic annotation and primer design for COI-based metabarcoding at the community level. This database integrates COI gene data from GenBank and mitochondrial genomes that are publicly available, followed by rigorous manual curation to eliminate redundancies and contaminants and to correct taxonomic annotations. We validate using the eKOI database for taxonomic annotation of protists by re-annotating several COI-based metabarcoding studies, revealing previously unidentified biodiversity. Phylogenetic analyses confirmed the accuracy of the taxonomic annotations, highlighting the potential of eKOI to uncover new biodiversity in various eukaryotic lineages.

## Introduction

Metabarcoding has become a powerful tool in the last two decades (Rondon et al., 2000), enabling researchers to comprehend biodiversity patterns without the biases inherent in traditional sampling methods (Taberlet et al., 2012). This is due to the ability to retrieve DNA directly from the environment (eDNA), allowing the characterization of microbial communities without the need for isolation or culture-dependent approaches (Ogram et al., 1987; Pawlowski et al., 2021). Furthermore, its low cost and ease of use have facilitated its application across diverse biological disciplines. For example, metabarcoding has provided novel insights into biogeographical (de Vargas et al., 2015; Vernette et al., 2021) and ecological (Burki et al., 2021; Deiner et al., 2017) patterns. Metabarcoding requires, however, well-curated and comprehensive reference taxonomic databases to provide accurate taxonomic assignments of the sequenced eDNA.

Ribosomal genes, like the 18S rRNA gene (18S), are the most widely used genetic markers for species delimitation and phylogenetic inference among eukaryotic microorganisms (protists). This is due to their universality (they are present in all living beings), the combined presence of conserved and hypervariable regions that provide a phylogenetic resolution at both high and low taxonomic levels, and the availability of different generalist and taxon-specific primer sets. As a result, several large taxonomic databases have been generated, such as PR^2^ (Guillou et al., 2013), which are essential for taxonomic annotation of metabarcoding studies.

However, the 18S presents constraints in its ability to resolve taxonomy at both the species and intraspecies levels, due to its highly conserved nature (Girard et al., 2022; Lara et al., 2022). To overcome this limitation, researchers have focused on more divergent regions within ribosomal genes, such as the internal transcribed spacer (ITS) (Op De Beeck et al., 2014), or rapidly evolving genes like the ribulose-bisphosphate carboxylase gene (rbcL) (Apothéloz-Perret-Gentil et al., 2021; Pichard et al., 1997). While these markers offer improved phylogenetic resolution for species delimitation, the taxonomic databases are biased towards specific groups, such as fungi or diatoms (Bell et al., 2017; Větrovský et al., 2020), which restricts their wider applicability in eukaryotes.

The mitochondrial cytochrome oxidase subunit I (COI) gene has been proposed as the barcode gene for species delimitation in metazoans (Hebert et al., 2003). The COI gene has also started to be applied to the taxonomy and systematics of protists, such as in testate amoebae (González-Miguéns, Soler-Zamora, Villar-Depablo, et al., 2022; Kosakyan et al., 2016), foraminifera (Girard, Langerak, et al., 2022), coccolithophores (Hagino et al., 2011) and diatoms (Moniz & Kaczmarska, 2009). This has led to the creation of several reference COI taxonomic databases, like BOLD (Ratnasingham & Hebert, 2007), CO-ARBitrator (Heller et al., 2018) or MIDORI2 (Leray et al., 2022). However, these databases are predominantly biased toward metazoans, lacking curated and redundant-free sequences for other taxonomic groups, as well as standardized taxonomic ranks widely used for eukaryotes (Burki et al. 2020). These restrictions impede the application of the COI gene in community-level studies of eukaryotes that include protists, and obstruct taxonomic integration across different molecular marker databases, such as the widely used PR^2^ for 18S rRNA gene.

To address these limitations, we present here a novel and curated eukaryote-wide (covering 80 phyla) database of the mitochondrial COI gene, named eKOI. This new reference database aims to address the limitations of existing COI taxonomic databases at the eukaryote community-level by developing a comprehensive, well-curated taxonomy-wide resource with a focus on protists, analogous to PR^2^. This will facilitate the taxonomic annotation of metabarcoding sequences at the eukaryote community-level, as well as comparisons among different taxonomic databases derived from other molecular markers. To create this dataset, we combined COI gene data extracted from GenBank and the complete COI gene obtained from publicly available mitochondrial genomes. A thorough and manual curation process was implemented to eliminate redundant sequences, identify potential contaminants, and correct taxonomic annotation errors while refining the taxonomy assignation for each sequence. The accuracy of intron detection for sequences extracted from mitochondrial genomes was verified. We evaluated this new database by taxonomically reannotating various COI-based metabarcoding studies, resulting in the identification of previously unidentified diversity, and a comparison with existing COI taxonomic databases. Finally, we further validated the new taxonomic annotations for these studies by constructing phylogenetic trees using sequences from the eKOI database, confirming the large amount of previously uncharacterized biodiversity.

## Materials and methods

### GenBank database sequences downloading and curation

To construct the eKOI taxonomic database, we initially retrieved the sequences from the mitochondrial gene “Cytochrome oxidase subunit I” (COI) from GenBank. We established keywords to search for each taxonomic group in the "NCBI taxonomy browser": “((“X”[Organism] OR “X”[All Fields]) AND co1[All Fields]) OR ((“X”[Organism] OR “X”[All Fields]) AND cox1[All Fields]) OR ((“X”[Organism] OR X[All Fields]) AND coi[All Fields]) OR ((“X”[Organism] OR X[All Fields]) AND cytochrome oxidase[All Fields] AND subunit[All Fields] AND 1[All Fields]) OR ((“X”[Organism] OR “X”[All Fields]) AND cytochrome oxidase[All Fields] AND subunit[All Fields]) OR ((“X”[Organism] OR “X”[All Fields]) AND coxi[All Fields])”, where “X” denotes the name of each major taxonomic group of “NCBI taxonomic browser” within “Eukaryota”. The files were downloaded in “INSDSeq XML” format, obtaining about 4 million sequences. The resultant sequences, grouped by taxonomic groups, principally phyla, were processed using a custom script (1_sequences_procesing.py in supplementary data). This script eliminated the sequences that were duplicates, smaller than 200 bp and larger than 3,000 bp. Next, to reduce the redundancy and the total number of sequences, clusters were created based on similarity percentages using vsearch ver. 2.14.1 (Rognes et al., 2016, p. 20), selecting a representative sequence for each cluster. The similarity percentage was established at 97%, except for Arthropoda, Chordata, and Mollusca for which it was set at 90%, aiming to balance the number of sequences per taxonomic group in the final eKOI database. This limits identifications to low taxonomic levels such as species or genus. If taxonomic identification below these levels is desired, it is recommended to enrich the eKOI database with sequences from the targeted taxonomic group. Chimeric sequence detection was performed using vsearch ver. 2.14.1 “*de novo*” (Rognes et al., 2016), removing the chimeric sequences. Lastly, a “fasta” file was generated for each taxonomic group containing the sequences with the taxonomy string defined in GenBank.

Alignments were generated for each taxonomic group, using MAFFT ver. 7.490 (Katoh et al., 2002), using default parameters. Finally, manual curation of the resulting sequences was performed using the software Geneious Prime (ver. 2019.0.4), removing divergent sequences that may be errors or taxonomic misclassifications.

### Mitochondrial genome database curation and integration with GenBank database

The resulting curated sequences retrieved from GenBank were combined with the mitochondrial genome of public databases, such as GenBank and Zenodo. The COI gene was extracted from complete mitochondrial genomes present in GenBank based on the sequence annotations. Some resulting sequences contained exons and introns. It has been demonstrated that some introns can result from nuclear pseudogenes (Andújar et al., 2021). Therefore, we first tested whether the introns were potential pseudogenes or chimeric sequences. To accomplish this, two datasetswere generated from the mitochondrial genomes: one with the entire COI gene including introns and exons, and another one containing only the coding region, the exons.

To further test the presence of introns in the mitochondrial genomes, polymerase chain reaction (PCR) amplifications were performed on two cultures of Choanoflagellata and Filasterea species. We used one culture of Stephanoecidae sp. (Choanoflagellata) (strain BEAP0094) and one culture of *Capsaspora owczarzaki* (Filasterea) (strain ATCC30864). In the case of Stephanoecidae sp., the mitochondrial genome of the Choanoflagellata *Monosiga brevicollis* has an intron within the region of the Folmer et al. (1994) primers LCO 1490 (5’ GGTCAACAAATCATAAAGATATTGG 3’) and HCO 2198 (5’ TAAACTTCAGGGTGACCAAAAAATCA 3’) (Burger et al., 2003); on the other hand, the mitochondrial genome of *C. owczarzaki* has an intron in the region of the primer LCO (Suga et al., 2013).

The following PCR protocol was followed in all cases: a final reaction volume of 20 μL containing 6 μL of distilled water, 12 μL MyTaq Red DNA polymerase Mix (BioLine), 1 μL of each primer (10 μM), and 2 μL of culture, without DNA extraction. The PCR was performed using the universal mitochondrial COI primer pair LCO 1490 (5’ GGTCAACAAATCATAAAGATATTGG 3’) and HCO 2198 (5’ TAAACTTCAGGGTGACCAAAAAATCA 3’) (Folmer et al., 1994), with the following PCR cycling profile: initial denaturation at 96 °C for 5 min, followed by 40 cycles at 94 °C for 15 s, 40 °C for 15 s, and 72 °C for 90 s and a final extension step at 72 °C for 10 min. After the amplification, 3 μL of the reaction was analysed by electrophoresis on a 1% agarose gel to verify fragment size and check for contaminations. Bands with the expected size were excised from the gel and stored at 4 °C. The samples were sequenced using Sanger dideoxy-technology in both directions by the company Eurofins Genomics (https://eurofinsgenomics.eu/). The resulting sequences were uploaded to GenBank with accessions PQ097056-PQ097057.

Alignments were performed using MAFFT ver. 7.490 (Katoh et al., 2002), using default parameters, aiming to determine if the sequences from the GenBank database or the newly amplified sequences contained the intron region. To achieve this, we used the curated sequences from GenBank, the new sequences from Stephanoecidae sp. and *C. owczarzaki*, and the two mitochondrial genome COI databases (with and without introns). The script 2_percentage_identity_graphic.py (available in the supplementary data) graphically represents the percentage of identity for each position in an alignment. Once the introns were confirmed to be potential contaminations, the curated GenBank sequences were combined with the coding regions of the COI gene extracted from the mitochondrial genomes (see results section “Testing the presence of introns within COI gene”). The final fasta files and alignments for each taxonomic group are available from the Supplementary data. Finally, the alignments without introns were used to curate the sequences with a wrong taxonomy following the “curation process” in EukRef (del Campo et al., 2018), generating phylogenetic reference trees and alignments per phylum.

### Taxonomy path curation

One of the limitations of current molecular databases for the taxonomic assignment of eDNA sequences is how to handle the variable taxonomic ranks across clades of eukaryotes. Even though higher taxonomic ranks, such as order or class, lack comparable evolutionary context, in terms of divergence time or evolutionary history in general, many computational tools require a fixed number of ranks across taxa.

To achieve this, the names of the sequences were extracted into a CSV file using the script (3_fasta_name_extraction.py in supplementary data). This file was manually curated to correct the taxonomy of each sequence. The aim was to ensure that all sequences have "homologous" taxonomic categories among them at each taxonomic level. We used the taxonomic levels proposed by PR^2^ version 5.0 (released in 2023) (Guillou et al., 2013), comprising nine levels: “Domain; Supergroup; Division; Subdivision; Class; Order; Family; Genus; Species”. We also generated another dataset, to which we added the level “phylum” between "subdivision" and "class" since phylum is one of the most widely used taxonomic ranks in metazoans. Once each CSV file was manually curated, the names of the sequences in the fasta file were substituted by the accession identifier unique to each sequence, using the script (4_fasta_name_substitution.py in supplementary data). The final eKOI taxonomic database can be available from the supplementary data (file “eKOI.fasta”).

### Testing the eKOI database accuracy and comparing with other taxonomic databases

To test the accuracy of the eKOI database, and its capacity to characterize new biodiversity, 15 metabarcoding studies based on COI were selected (see Tables 1 and 2 for details on metabarcoding studies and primers used). The raw data of these studies was re-analysed using the protocol described in (González-Miguéns et al., 2023), using the dada2 R package (Callahan et al., 2016). Subsequently, the resulting amplicon sequence variants (ASVs) smaller than 100 bp were discarded, as taxonomic assignments for fragments of such small size are typically unreliable.

**Table 1.**
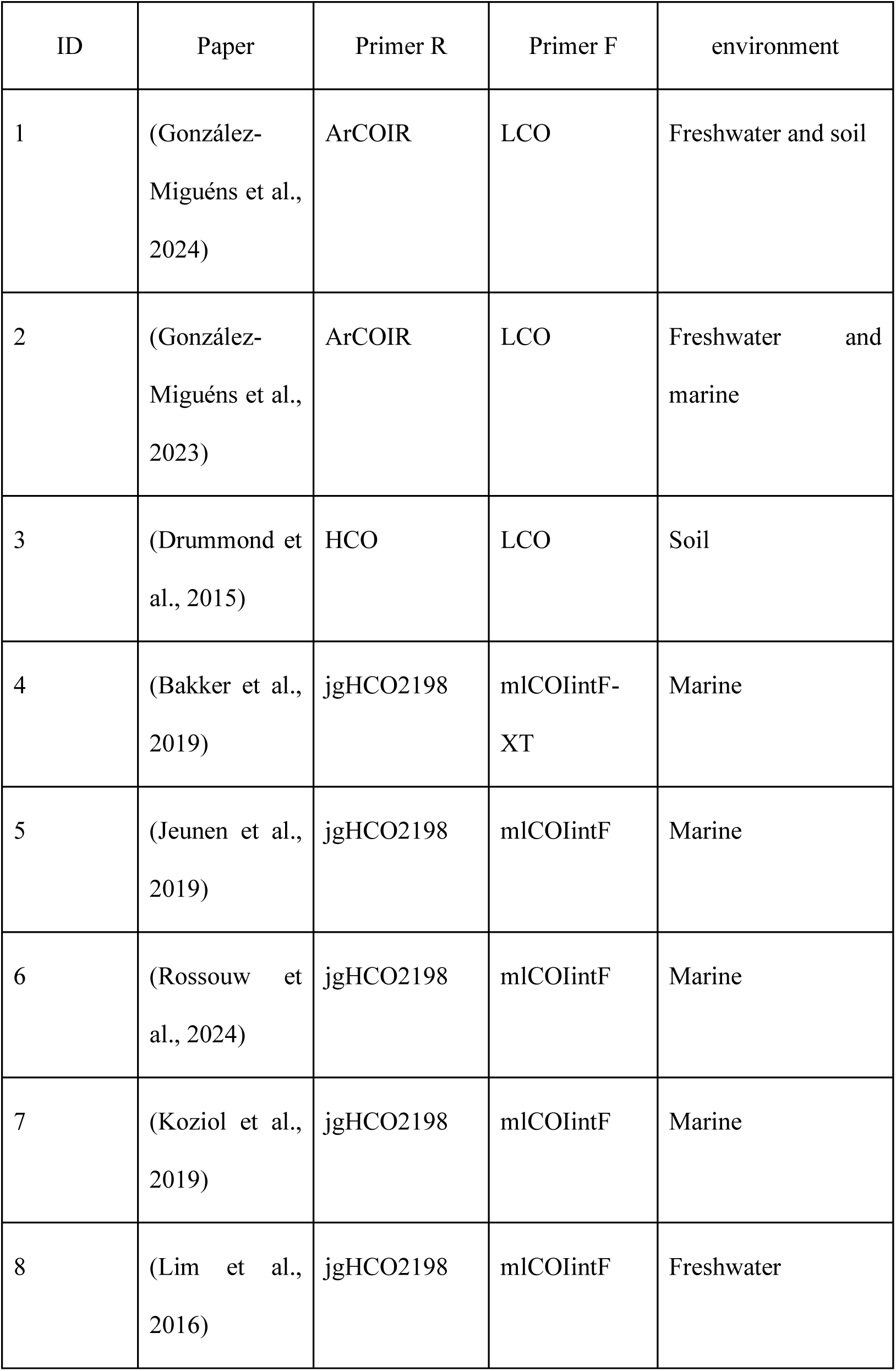
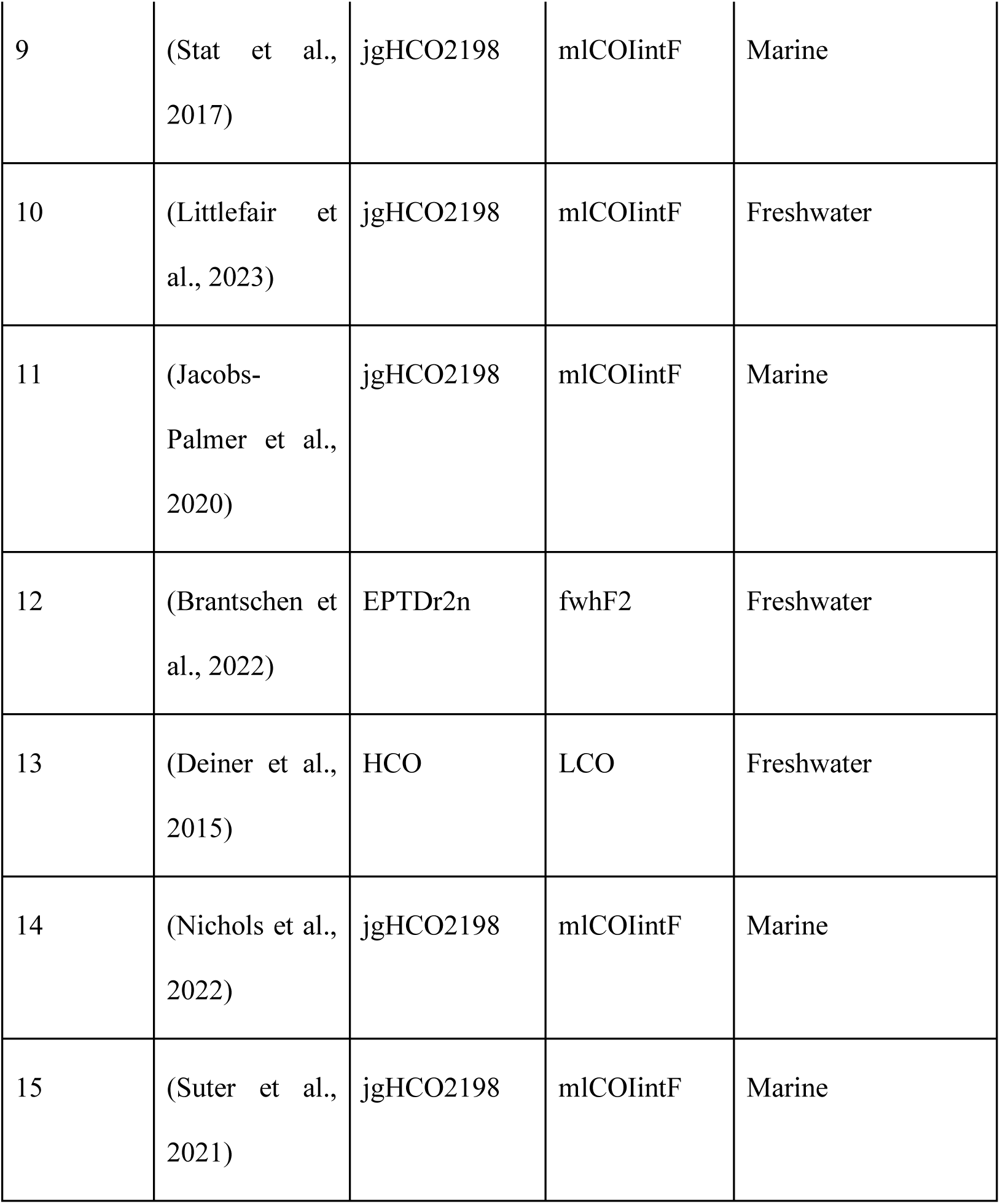
Metabarcoding studies based on the COI molecular marker that has been re-analyzed with our new eKOI dataset, together with the primers used in each case and the environment.

**Table 2.**
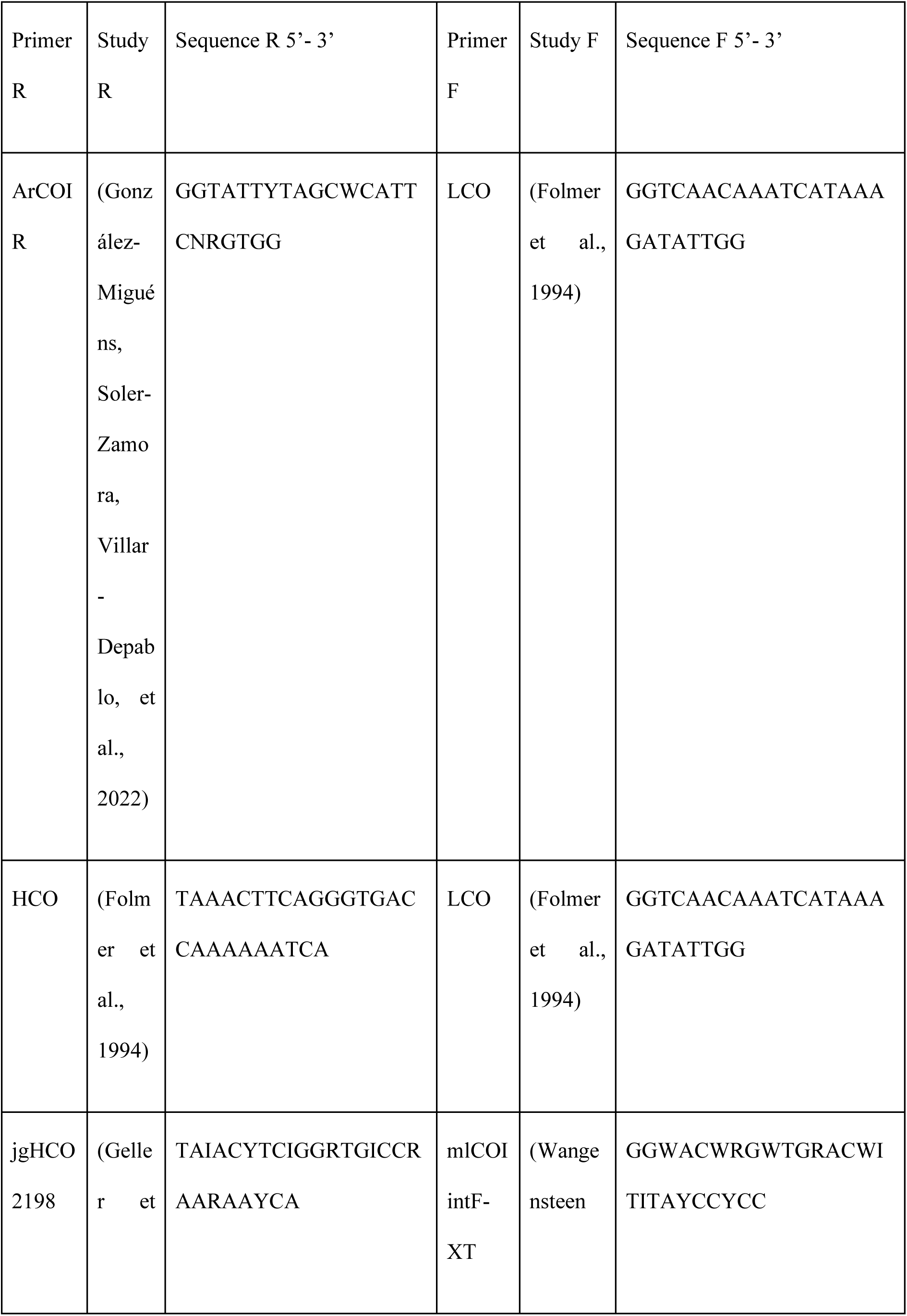
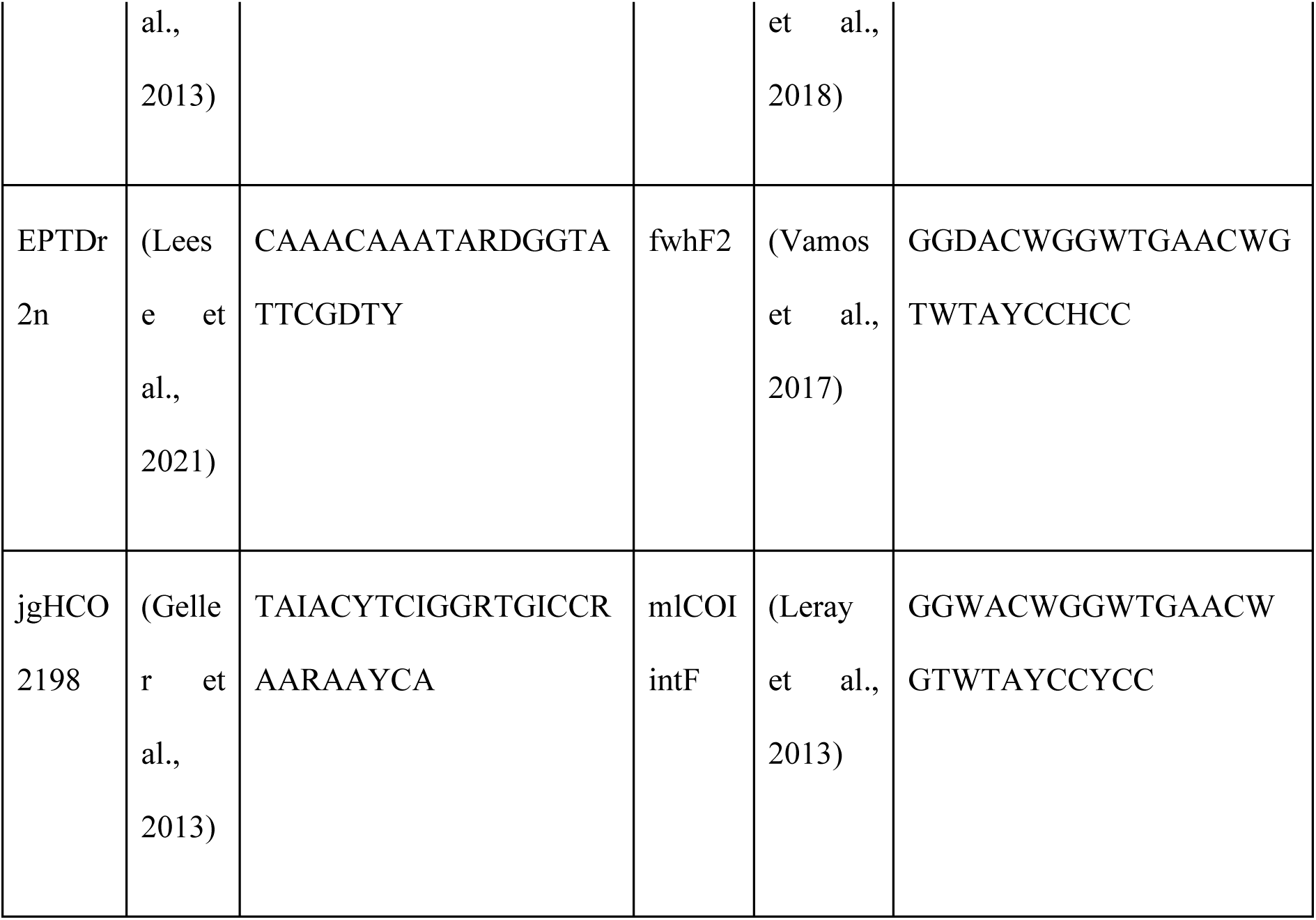
Combinations of forward (F) and reverse (R) primers.

Taxonomic assignment was performed using the eKOI and MIDORI2 (Leray et al., 2022) databases. For this purpose, a custom script (5_taxonomic_assignation.py in supplementary data) was employed. This script generates an independent folder for each fasta file present in the script’s directory. Within each folder, an Excel file is generated containing the taxonomic assignment information for each ASV using vsearch usearch_global command (ver. v2.14.1, Rognes et al., 2016). ASVs with lower than 84% similarity to reference sequences were not considered. This threshold was established based on the order Arcellinida and metazoans COI pairwise distance (González-Miguéns et al., 2023; Hebert et al., 2003). Subsequently, a fasta file is generated for each desired taxonomic level. In this case, we selected phylum. The resulting ASVs for each group taxonomically assigned with eKOI can be downloaded from “3_ASV_metabarcoding” in the supplementary data.

To graphically represent the diversity of each taxonomic group and test the accuracy of taxonomic assignments using eKOI, we generated phylogenetic trees for each taxonomic group, focusing on protists. To reduce the number of sequences for graphical representation, the ASVs were grouped into Operational Taxonomic Units (OTUs) based on a similarity percentage of 97% using a custom script (6_cluster_OTU.py in supplementary data). Alignments of the resulting OTUs were performed using MAFFT ver. 7.490 (Katoh et al., 2002), with default parameters, along with sequences from the eKOI database of the same taxonomic groups, and at least two sequences from sister groups as outgroups. Tree topologies and node supports were evaluated using maximum likelihood (ML) with IQTREE2 version 2.0, where the best substitution models were selected with ModelFinder (Kalyaanamoorthy et al., 2017). Node supports were assessed with 10,000 ultrafast bootstrap replicate approximations (Hoang et al., 2018). The resulting trees were graphically edited using the R package ggtree (Yu et al., 2017), highlighting the ecology of each generated OTU.

For some groups (Apicomplexa, Cercozoa, Filasterea, and Heterolobosea), reconstructions of ancestral ecology characters were performed to illustrate the potential of the eKOI database. For the reconstruction of ancestral habitat characters, we used the phytools R package (Revell, 2012). We employed the function *make.simmap* (Bollback, 2006) to generate 500 stochastic character maps from our dataset, under the equal rates model. These sets of stochastic maps were then summarized using the function *densityMap*, plotting the posterior probability of being in each state across all the edges and nodes of the tree. For the graphical representation of the distribution maps of the new eDNA sequences, we used the maps R package.

Once taxonomic assignments were made, the different samples were grouped by study (Table 1) and then by ecology. From these groups, the mean ASV taxonomic assignment values were obtained across all samples using the script (7_taxonomic_assignation_mean.py in supplementary data). This script generates a CSV with the percentage of the number of ASV assigned to each phylum. This was performed for the eKOI and MIDORI2 databases (“5_COI_databses_comparison” in supplementary data). Boxplots were generated in R using ggplot2, selecting the phyla with at least 1% of ASV taxonomically assigned in each study.

## Results

### eKOI database

The final eKOI database includes 15,947 sequences, representing 80 phyla, 231 classes, 796 orders, and 2,646 families of eukaryotes (Fig. 1 and “eKOI_taxonomy.csv” in supplementary data). Therefore, eKOI expands the coverage of protist taxonomic groups compared to other COI databases, like MIDORI2 and incorporates eukaryotic groups that are currently absent in existing databases, such as Picozoa, Nibbleridia, or Rozellomycota (see Tables S1 and S2 in supplementary data). Almost every eukaryotic phylum in the eKOI database is represented by at least one complete COI gene sequence. The eKOI database predominantly contains sequences of two lengths: approximately 600 base pairs and 1,600 base pairs (Fig. 1). The ∼600 bp sequences correspond to fragments amplified using the HCO and LCO primer pair (Folmer et al., 1994), while the ∼1,600 bp sequences represent the complete COI gene obtained from mitochondrial genomes. The size of the coding part of the COI gene (exon, excluding introns) remains relatively stable across different eukaryotic phyla.

**Figure 1.**
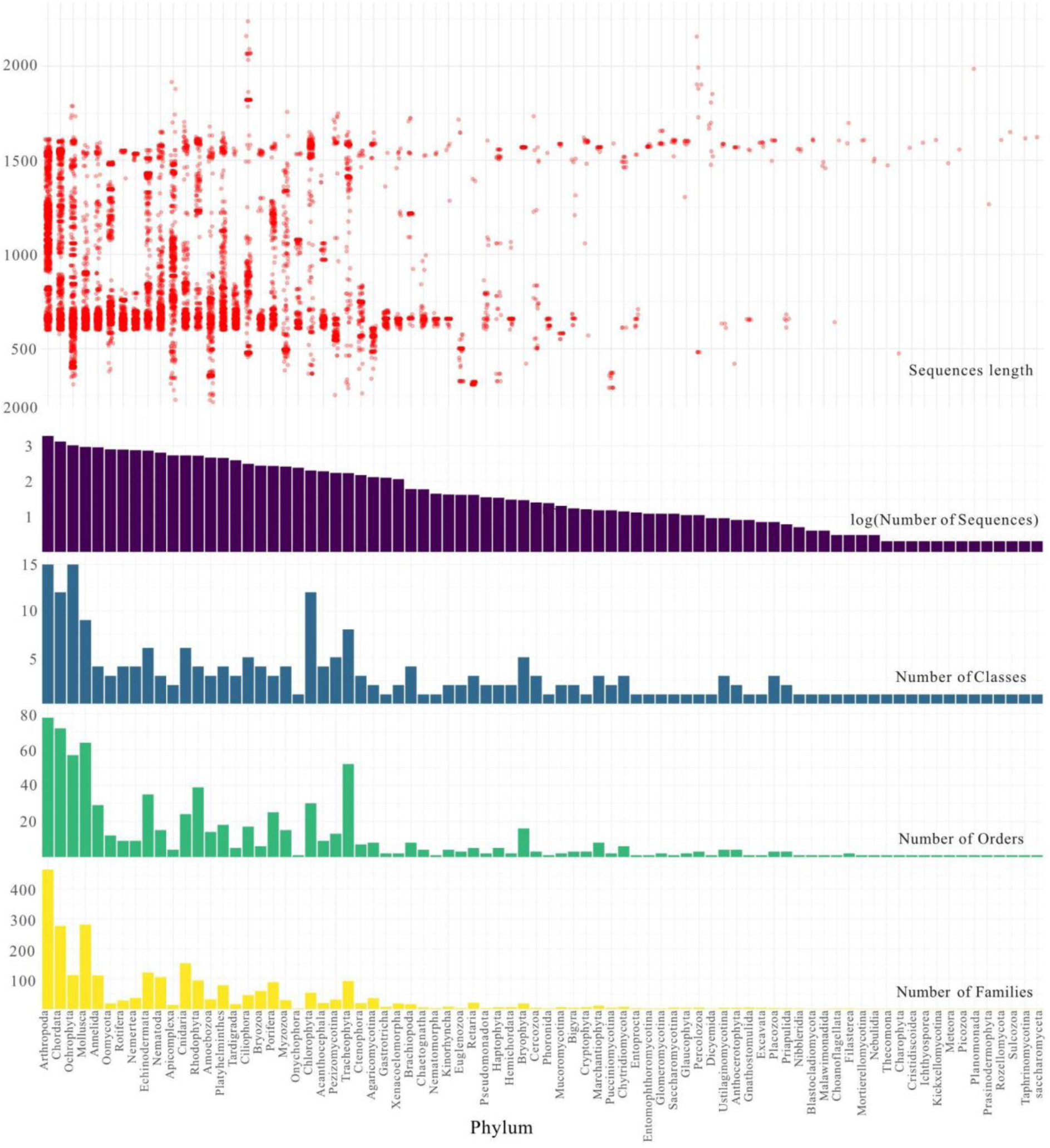
Graphs representing the number of families (yellow), orders (green), classes (blue), and total sequences (violet) per phylum present in the eKOI database. Red circles represent the size, in base pairs, of each sequence included in each phylum.

To validate the taxonomic annotations of the sequences retrieved and the presence of introns in the mitochondrial COI gene, two alignments were built, one including introns and another excluding them (see Materials and Methods, “Mitochondrial Genome Database Curation and Integration with GenBank Database”). None of the sequences obtained from GenBank COI sequences had introns (supplementary data folder “2_alignament_eKOI”). In the alignment including introns, GenBank COI sequences did not contain introns, resulting in large gaps within these sequences (Fig. S2). Furthermore, PCR amplifications were conducted on Stephanoecidae sp. (Choanoflagellata) and *Capsaspora owczarzaki* (Filasterea), targeting the regions where introns were previously annotated in the COI gene of the mitochondrial genomes, but no introns were detected in the amplified sequences from these regions.. The absence of highly divergent sequences in the alignments, as well as the lack of long branches in the phylogenetic trees, confirms the taxonomy of the sequences included in the eKOI database (supplementary data folder “2_alignament_eKOI”).

### Applying the eKOI database in metabarcoding studies

To test the effectiveness of the new eKOI dataset to taxonomically assign eukaryotic COI eDNA diversity, we applied it to fifteen metabarcoding studies (Table 1 and 2 and Fig. 2). The extensive use of primers jgHCO2198 (Geller et al., 2013), mlCOIintF (Leray et al., 2013) and mlCOIintF-XT (Wangensteen et al., 2018) is attributed to their ability to amplify a diverse range of eukaryotic taxonomic groups. The variation in the proportion of phyla within similar ecosystem types, using identical primers, is linked to differences in sample substrates, such as pelagic and plankton (Bakker et al., 2019), plankton (Suter et al., 2021), or sediments (Koziol et al., 2019). Across all studies, a significant diversity of eukaryotic microorganisms, such as Amoebozoa, Chlorophyta, Choanoflagellata, and Picozoa, was identified using eKOI database, constituting a substantial portion of all ASVs from each study (Fig. 2 and “eKOI_MIDORI2_comparison.xlsx” in supplementary data). Notably, the diversity within these taxonomic groups was high, revealing substantial underestimation of protist diversity and highlighting previously overlooked taxonomic groups in COI metabarcoding studies, such as choanoflagellates and Picozoa (Fig. 3 and S3), that were not recovered using MIDORI2 (Fig. 2 and “eKOI_MIDORI2_comparison.xlsx” in supplementary data). Instead, the use of the MIDORI2 database results in a higher representation of metazoans compared to other eukaryotic groups, in contrast to the assignments provided by eKOI (Fig. S4).

**Figure 2.**
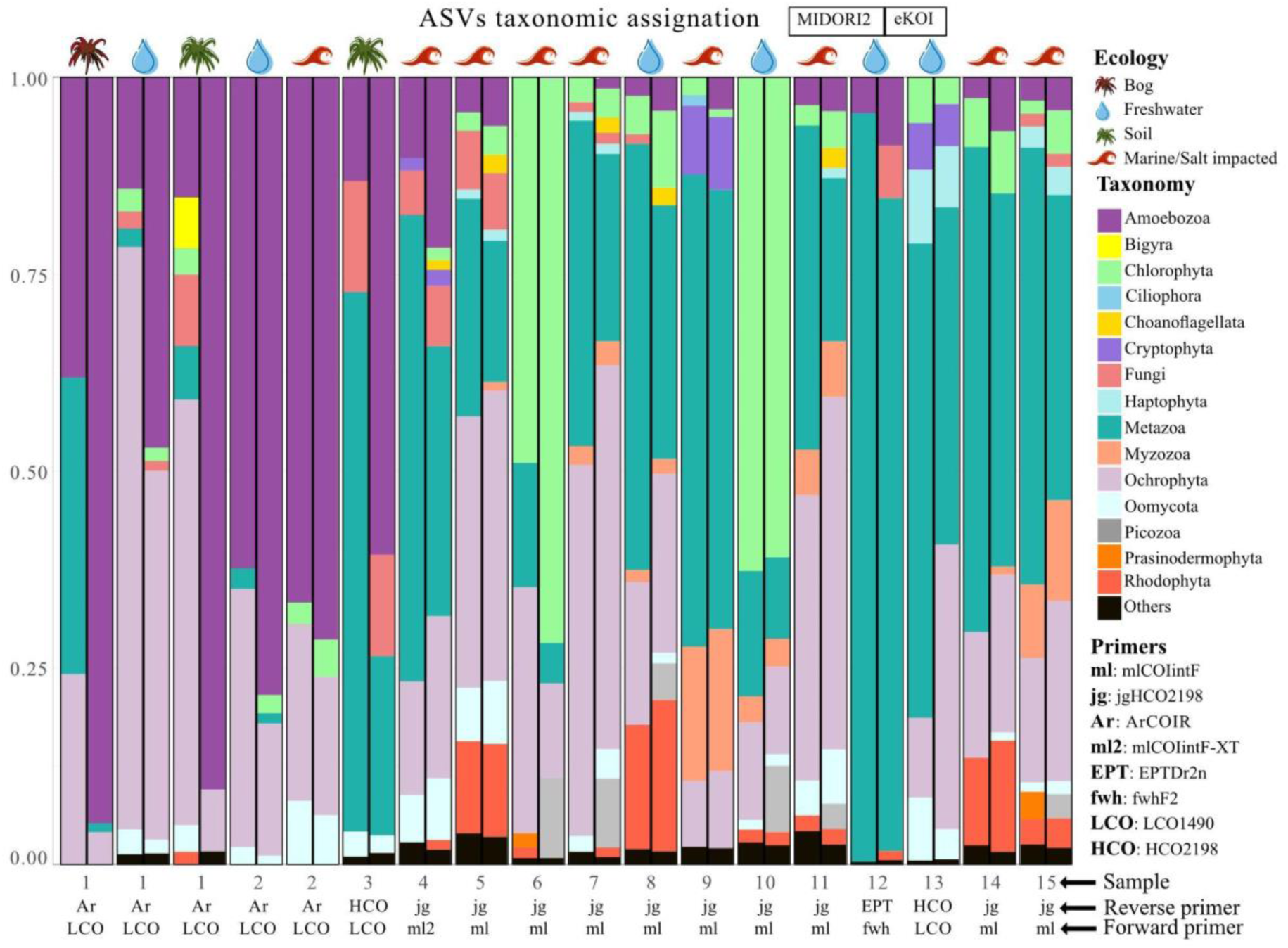
Bar chart representing the proportion of ASVs, relative to the total taxonomically classified, to each phylum in the MIDORI2 (left boxplots) eKOI database (right boxplots) from 15 metabarcoding studies (Table 1). Due to the large number of phyla present in each study, only the phyla with at least 1% of ratio per study are represented (the rest of the other phyla are represented as “others”). The metazoan phyla were grouped as "Metazoa". The environment type of each study is represented by different symbols. The pair of primers used in each study is also presented.

**Figure 3.**
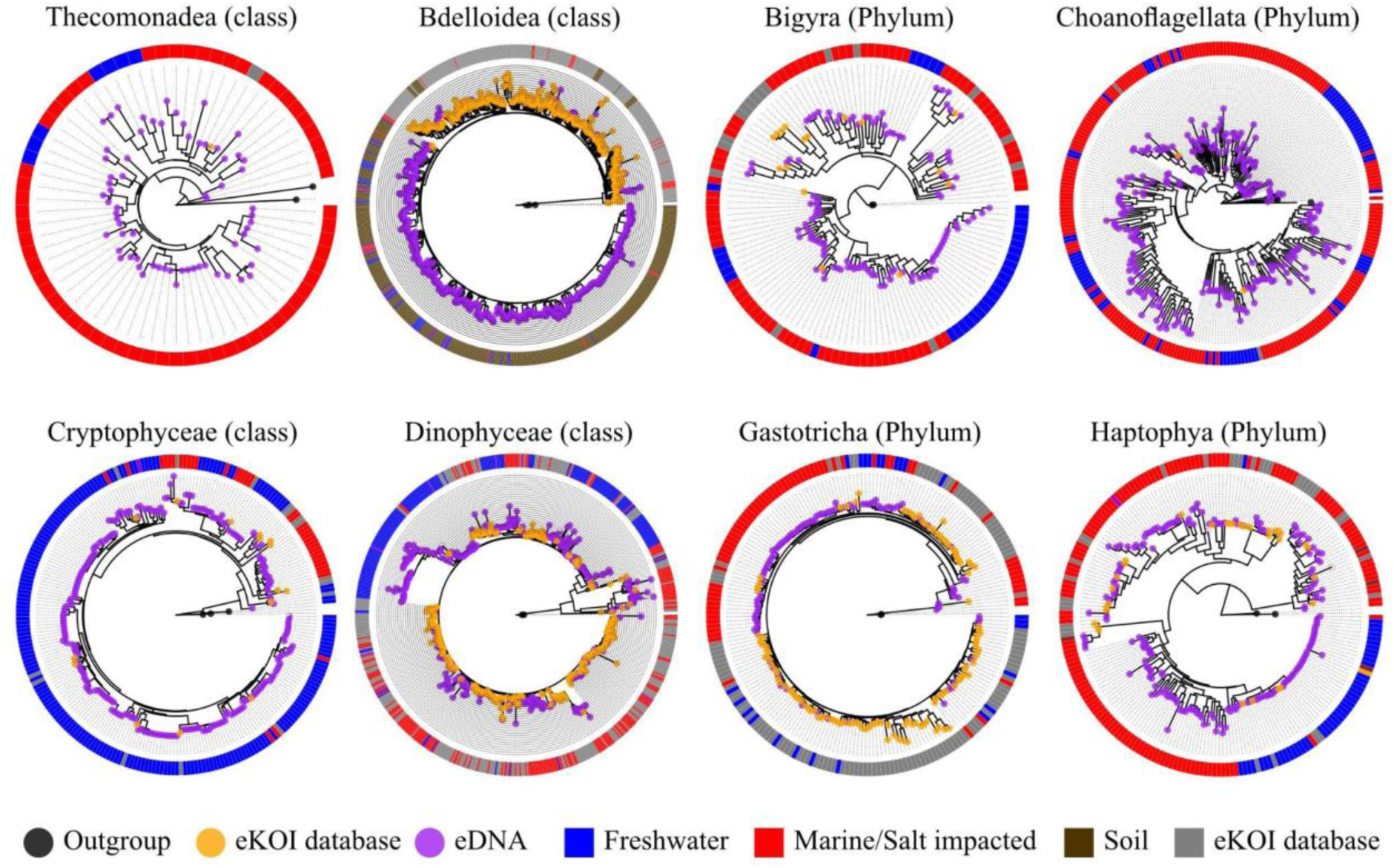
Phylogenetic trees obtained by combining sequences from the eKOI database of different taxonomic groups (orange circle), with outgroups (grey circle) and eDNA OTUs (purple circle). OTUs were obtained from similarity clustering of taxonomically reannotated ASVs from metabarcoding studies (Table 1). The circle surrounding each phylogenetic tree represents the environment of each OTU, sequences from the eKOI database are represented in grey, with the ecology not represented to reflect the newly characterised biodiversity for each taxonomic group.

The utility of ASVs derived from eDNA samples extends beyond characterizing novel molecular biodiversity. For instance, we examined the Cyphoderiidae (Rhizaria), a group in which ecological transitions across the salinity barrier have influenced its evolutionary history (González-Miguéns, Soler-Zamora, Useros, et al., 2022). While these transitions were primarily characterized using the 18S marker, the COI database lacked marine lineages that were exclusively obtained with the nuclear marker. The integration of new COI ASVs (Table 1), taxonomically classified as Cyphoderiidae using the eKOI database, enabled the characterization of marine lineages that had been previously identified with 18S, like the “marine clade 1” (González-Miguéns, Soler-Zamora, Useros, et al., 2022). This increase in the Cyphoderiidae COI sequence database enhancesthe robustness of the characterization of ecological transitions from marine to freshwater environments in Cyphoderiidae (Fig. 4), allowing the recovery of consistent ecological and phylogenetic patterns using both nuclear and mitochondrial molecular markers. This approach can be applied to other difficult-to-sample microorganisms, such as free-living or parasitic taxonomic groups; for example, Conoidasida (Apicomplexa) and Vahlkampfiidae (Heterolobosea) (Fig. 4).

**Figure 4.**
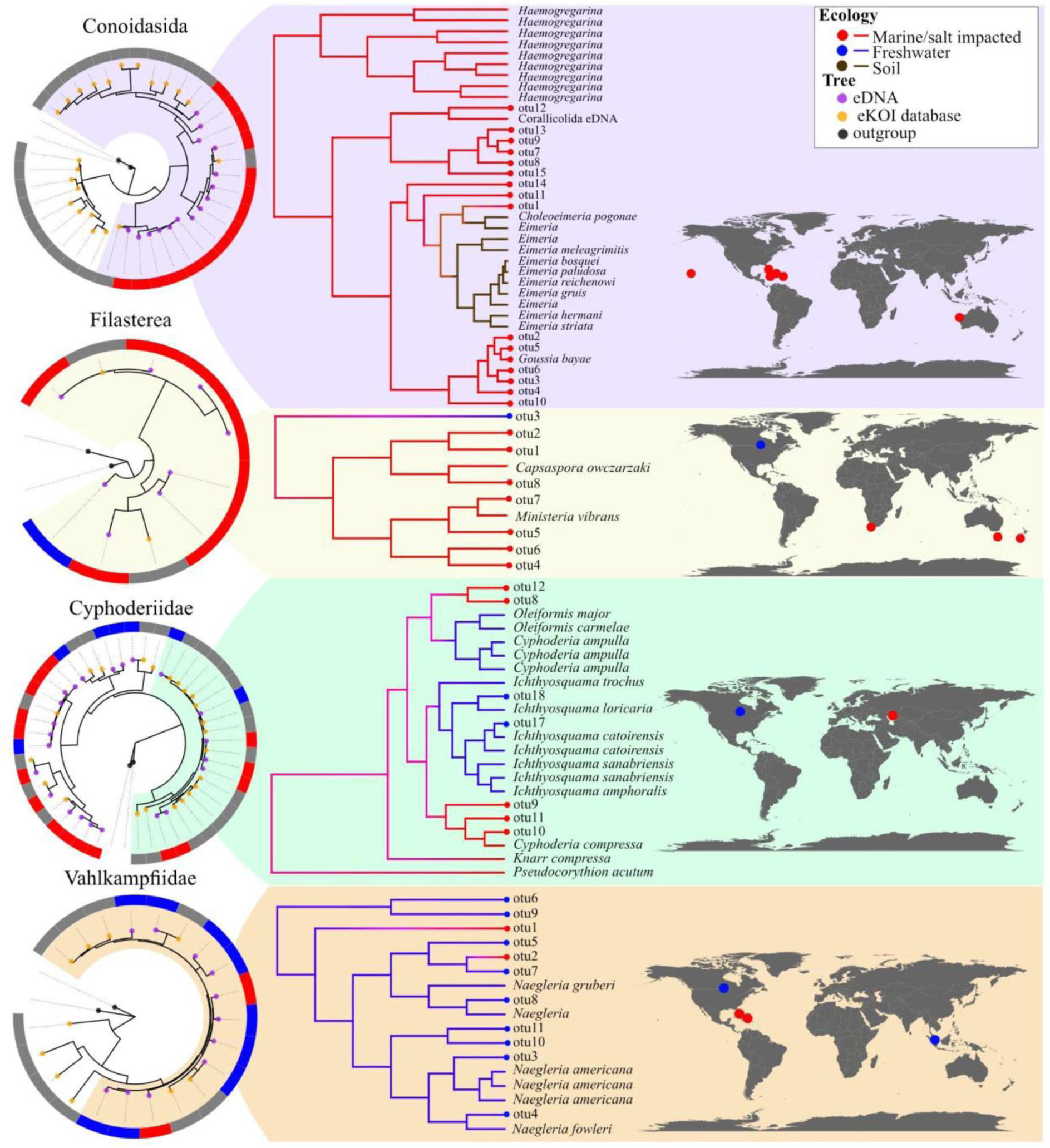
Phylogenetic trees obtained by combining sequences from the eKOI database of each taxonomic group (orange circle) and with outgroups (grey circle). The eDNA OTUs (purple circle) were calculated and generated independently for each taxonomic group, with the IDs shared across groups. The circle surrounding each phylogenetic tree represents the habitat of each OTU, sequences from the eKOI database are represented in grey, to reflect the newly characterised biodiversity for each taxonomic group. From each taxonomic group, a part of the tree was selected from which the ancestral habitat was reconstructed, as an example. We also represented with dots on the map (blue freshwater, brown soil, and red marine) the origin of the different OTUs were recovered.

## Discussion

The novel eKOI database presented here enables precise taxonomic assignments of sequences originating from eukaryotic COI metabarcoding studies, in particular those focusing on protists. The inclusion of curated protistsequences mitigates certain limitations present in existing COI databases focused on metazoans. Reducing redundant sequences improves the application of eKOI in community-level studies of eukaryotes; however, this limits its utility for species-level assignments of metazoans. For these specific assignments, there are databases specialized in metazoans, such as BOLD (Ratnasingham & Hebert, 2013) or MIDORI2 (Leray et al., 2022)as well as others focused on specific taxonomic taxonomic groups, like insects (Magoga et al., 2022), metazoans (Heller et al., 2018) or zooplankton (Bucklin et al., 2021).

The taxonomic curation of sequences in eKOI, along with their standardization with other sequence databases such as PR^2^, which focuses on the 18S ribosomal gene, enables comparison of taxonomic assignment results. Currently, most metabarcoding studies targeting the community level (Oliverio et al., 2020) or specific protist groups rely on the 18S rRNA gene (Segawa et al., 2018). However, a notable limitation of this marker is the phylogenetic resolution at lower taxonomic levels, due to its slow mutation rate compared to mitochondrial genes. While these traits are valuable to inferring relationships among “deep” taxonomic levels, their utility diminishes when attempting to discern relationships at the species or intraspecific level (González-Miguéns et al., 2023; Lara et al., 2022), with some exceptions (Ribeiro et al., 2024; Tragin & Vaulot, 2019). To solve this problem, fast-evolving genes, such as COI, are often used, providing species-level resolution (Kosakyan et al., 2016; Pinseel et al., 2020). In contrast, characterizing new divergent lineages of eukaryotes still presents challenges with COI due to its rapid mutation rate, which results in high divergence between the sequences of newly described lineages and those of known ones in established databases. The eKOI database comprises sequences for the entire COI gene for nearly all known eukaryotic phyla (Fig. 1), but lacks some phyla, like Dimorpha, Hemimastigophora, or Telonemia. Therefore, we propose that combining these independent nuclear and mitochondrial molecular markers could be ideal to uncover hidden patterns that may not be detected when relying on a single marker alone. Integrating mitochondrial eKOI (COI) and nuclear PR^2^ (18S) databases will provide new perspectives for hypothesis testing using eDNA.

Another aim of eKOI database is to facilitate the development of taxonomic-specific primers, similar to the existing applications for 18S like pr2-primers (Vaulot et al., 2022) that leverage the PR^2^ database to design primers for targeted taxonomic groups. This will allow applications of metabarcoding to reach beyond community-level analyses, with specific protocols targeting specific taxonomic groups. Two examples are the use of COI metabarcoding for Arcellinida (González-Miguéns et al., 2023) and foraminifera (Girard, Langerak, et al., 2022). This enables applied studies, such as the development of bioindicators (González-Miguéns et al., 2024) and ecological assessments (Girard, Macher, et al., 2022), to be focused on specific taxonomic groups using the COI gene. Furthermore, the integration of the sequences in the eKOI database with eDNA-derived ASVs could offer a valuable tool for inferring their biogeographical, diversification, or ecological patterns in future metabarcoding studies. This is exemplified in Fig. 4, although the number of samples and studies used is currently too small to infer patterns. These examples illustrate the potential of developing specific metabarcoding protocols targeting under-studied protist groups, often hindered by sampling difficulties or the impossibility to culture some lineages.

## Acknowledgments

This project has been funded by the European Research Council (ERC, MISSINGRELATIVES, 101097659) and the European Union’s Horizon 2020 research and innovation programme (grant agreement No. 949745). Views and opinions expressed are however those of the authors only and do not necessarily reflect those of the European Union or the European Research Council Executive Agency. Neither the European Union nor the granting authority can be held responsible for them. We zenoalso acknowledge support from the Departament de Recerca i Universitats de la Generalitat de Catalunya (exp. 2021 SGR 00751) and by PIE-202120E047- Conexiones-Life.

## Conflict of interest statement

The authors declare no competing interests.

## Data Availability Statement

The raw data, such as alignments and phylogenetic trees generated in this study have been deposited in GitHub https://github.com/rubenmiguens/eKOI_taxonomy_database.git, in the supplementary materials. The resulting eKOI taxonomic database can be downloaded from the supplementary data, Zenodo 10.5281/zenodo.12807325 (pending article acceptance) and PR^2^ webpage (pending article acceptance). The sequences obtained were uploaded to GenBank with accessions PQ097056-PQ097057.

---

**Figure S1.**
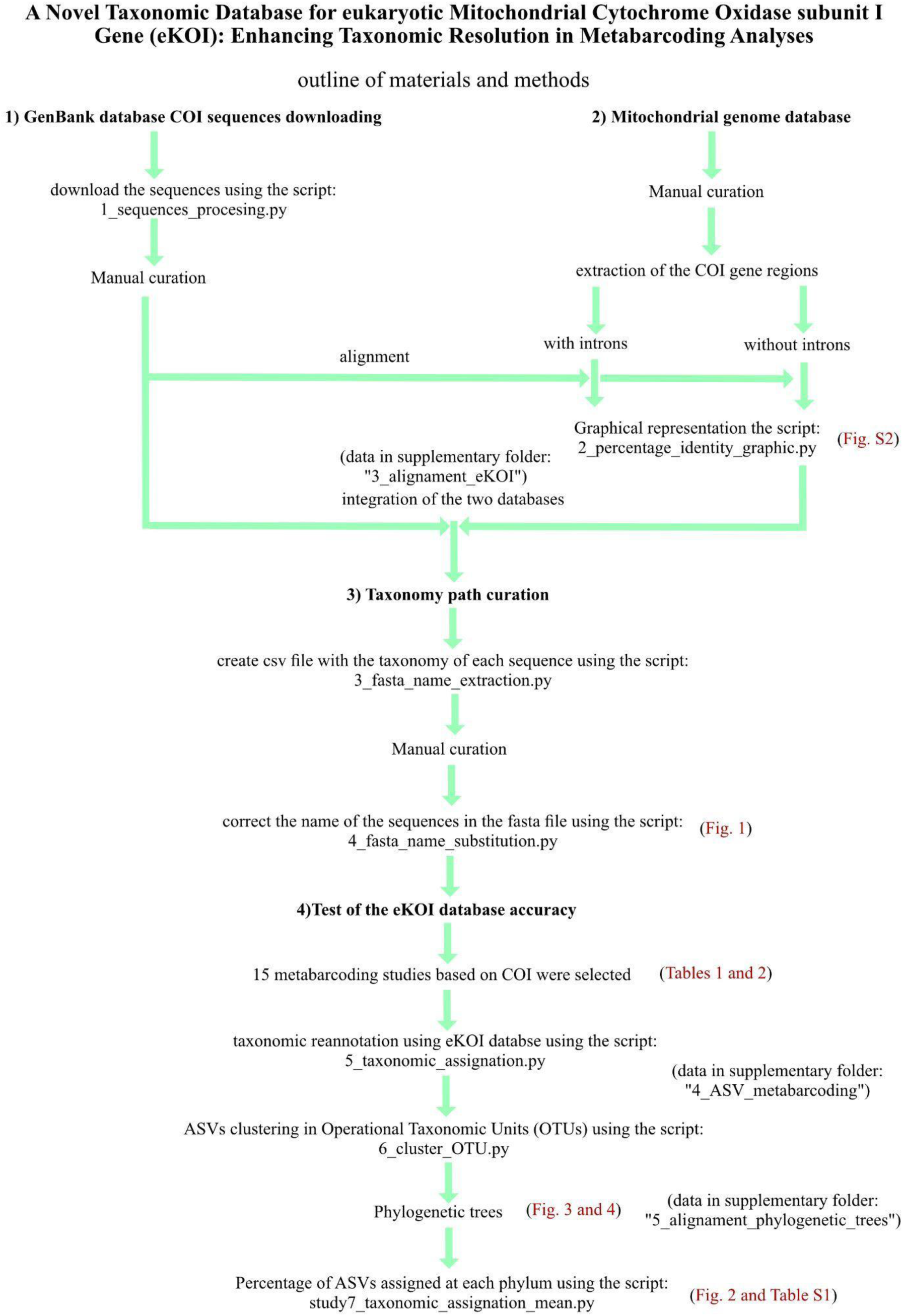
Schematic representation of the points covered through the materials and methods used in the study.

**Figure S2.**
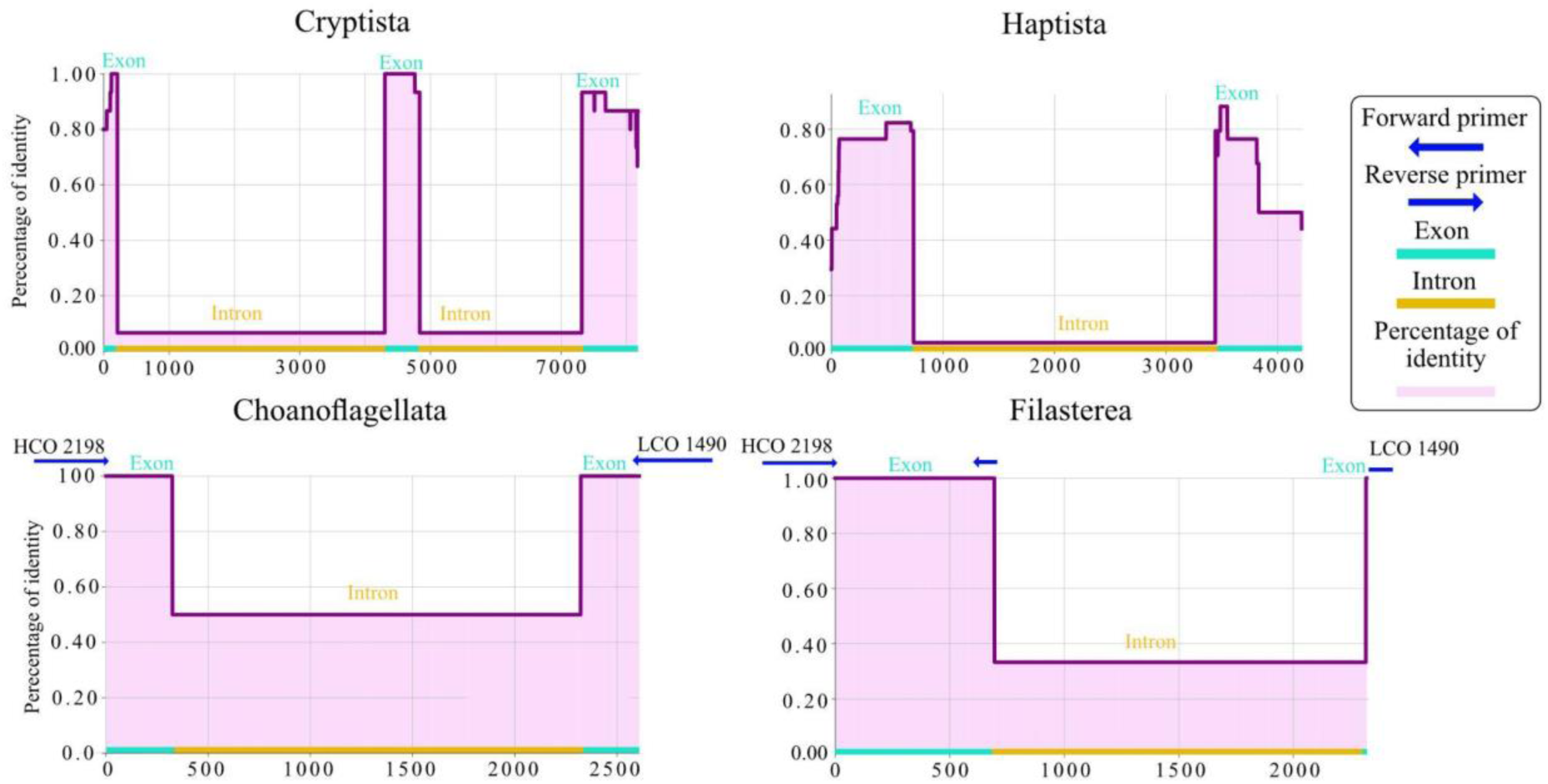
Plots representing the percentage identity by base, in purple, of the alignments generated from GenBank sequences with the extraction of the COI gene from the mitochondrial genomes. Regions characterised as exons and introns in the mitochondrial genomes are shown in blue and introns in yellow. The blue arrows represent the region, in the mitochondrial genomes, where the HCO and LCO primers used to generate *de novo* sequences from cultures are attached.

**Figure S3.**
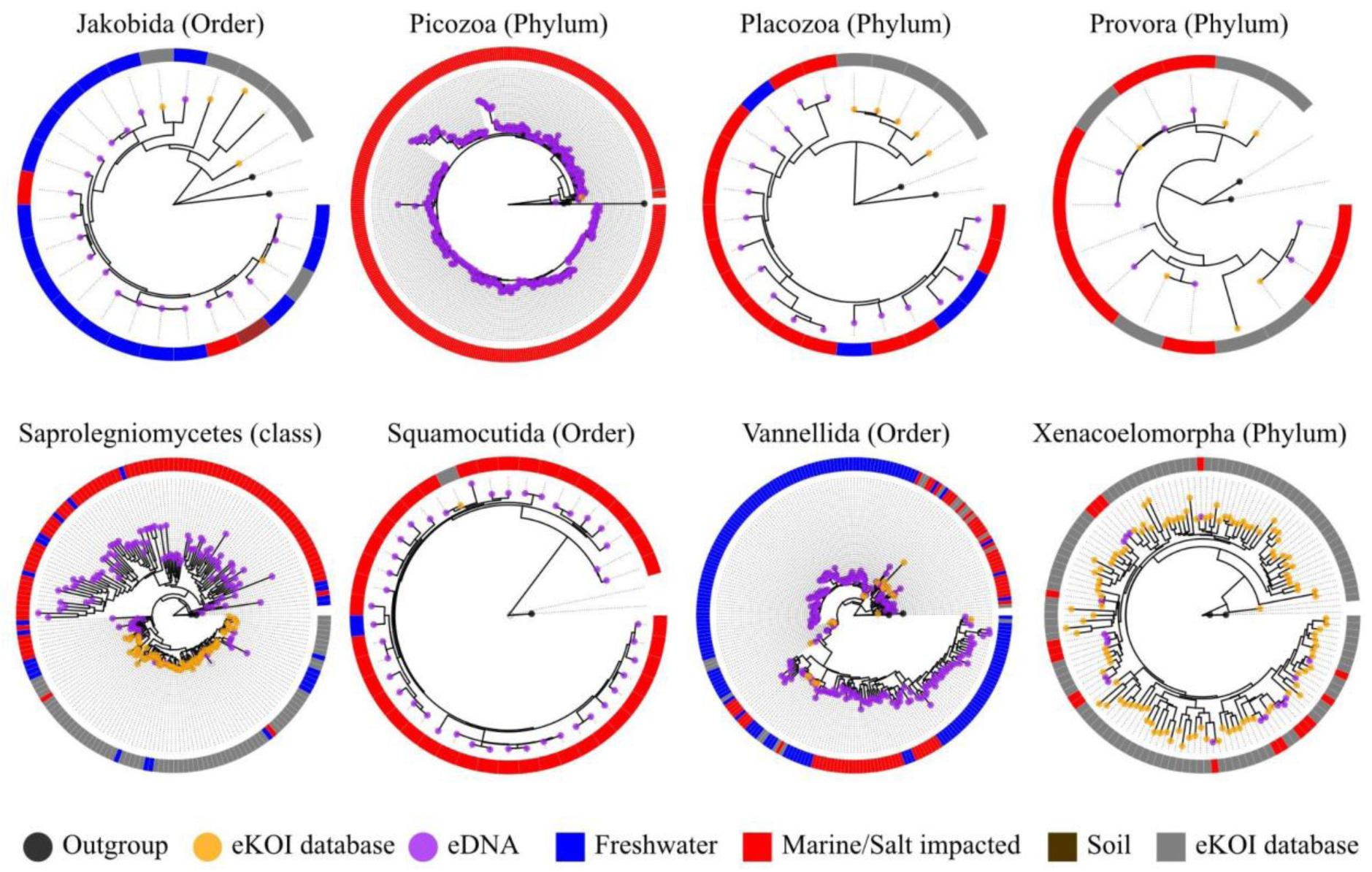
Phylogenetic trees obtained by combining sequences from the eKOI database of different taxonomic groups (orange circle), with outgroups (grey circle) and eDNA OTUs (purple circle). OTUs were obtained from similarity clustering of taxonomically reannotated ASVs from metabarcoding studies (Table 1). The circle surrounding each phylogenetic tree represents the environment of each OTU, sequences from the eKOI database are represented in grey, with the ecology not represented to reflect the newly characterised biodiversity for each taxonomic group.

**Figure S4.**
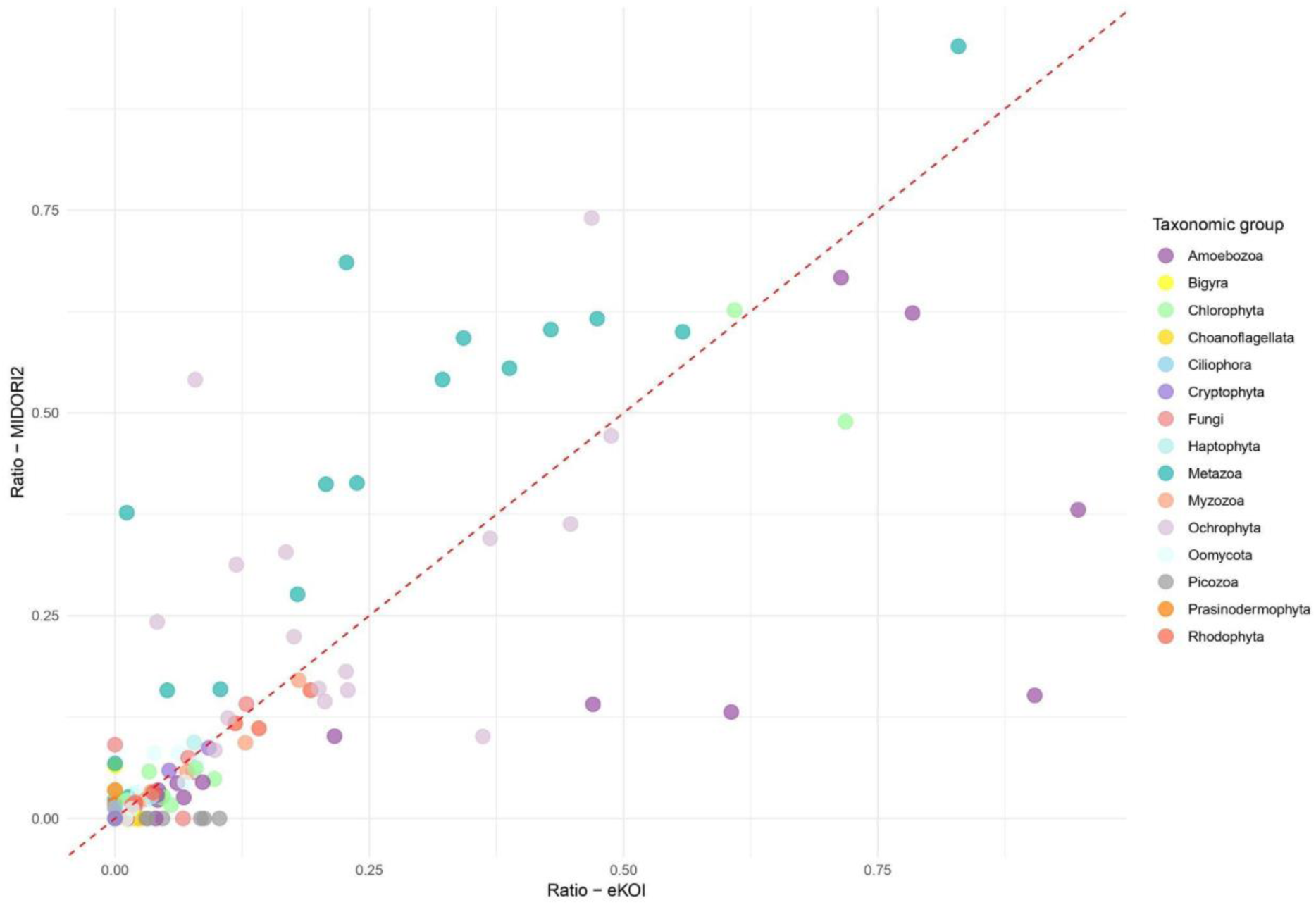
The plot compares the proportions of taxonomic groups between MIDORI2 and eKOI databases. Each point represents a taxonomic group, with colors distinguishing different organisms. The dashed red line indicates a perfect 1:1 ratio between the databases. Points on axes represent taxa present in one database but absent in the other (ratio = 0).

## Notes

https://github.com/rubenmiguens/eKOI_taxonomy_database.git

### Competing Interest Statement

The authors have declared no competing interest.

