## Supplementary figures and images for "A Novel Taxonomic Database for eukaryotic Mitochondrial Cytochrome Oxidase subunit I Gene (eKOI): Enhancing taxonomic resolution at community-level in metabarcoding analyses"

### S1

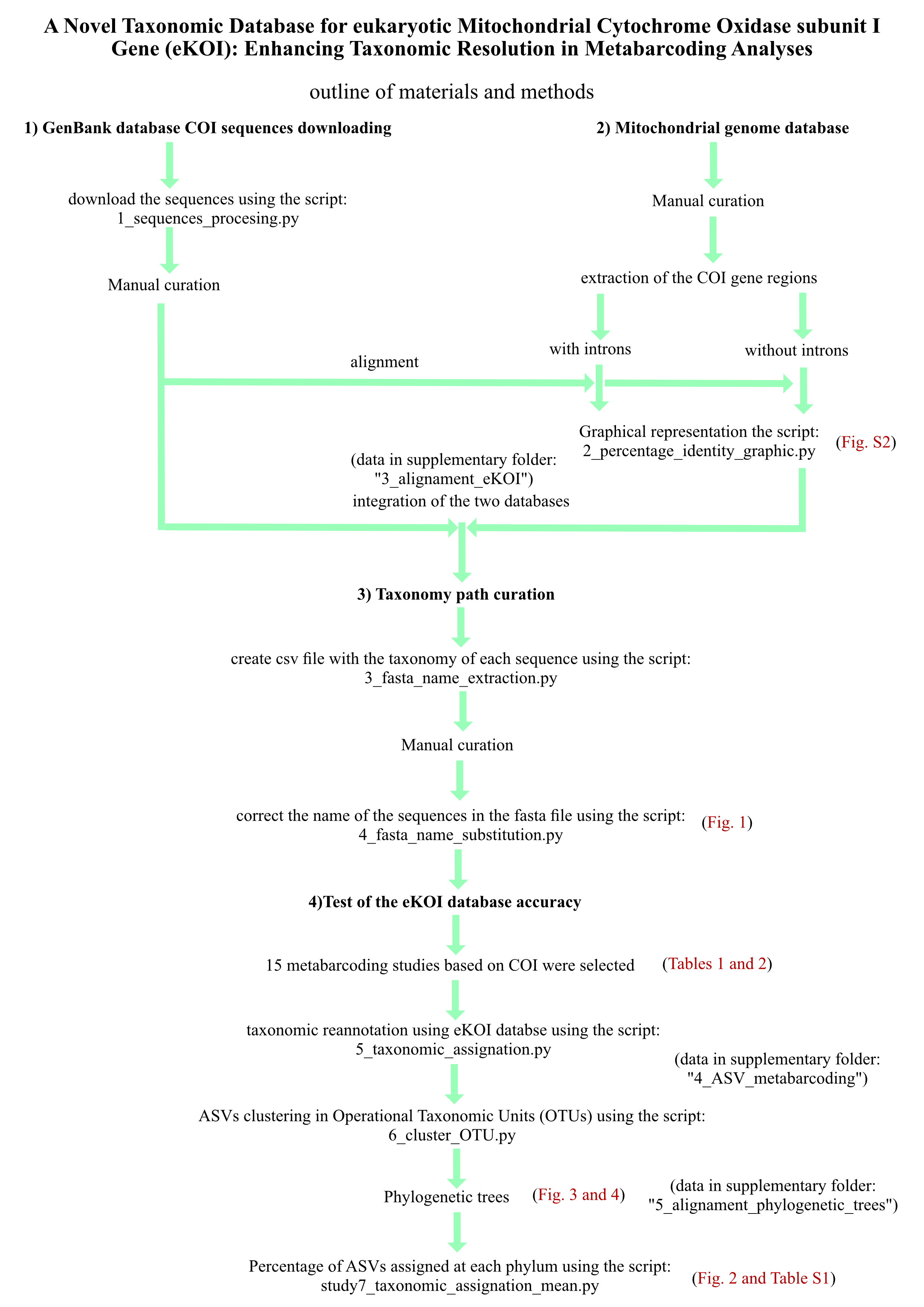

### S2

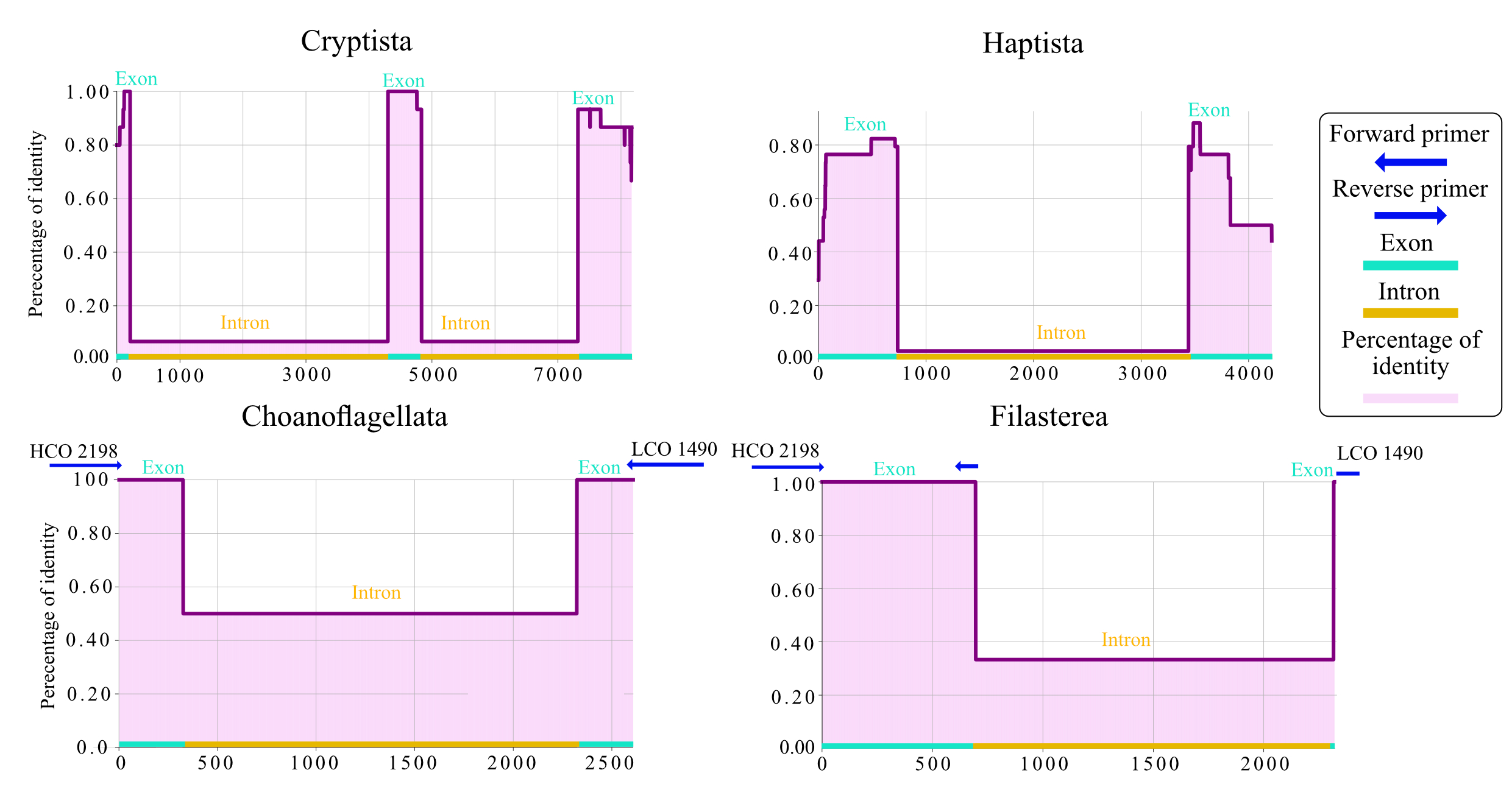

### S3

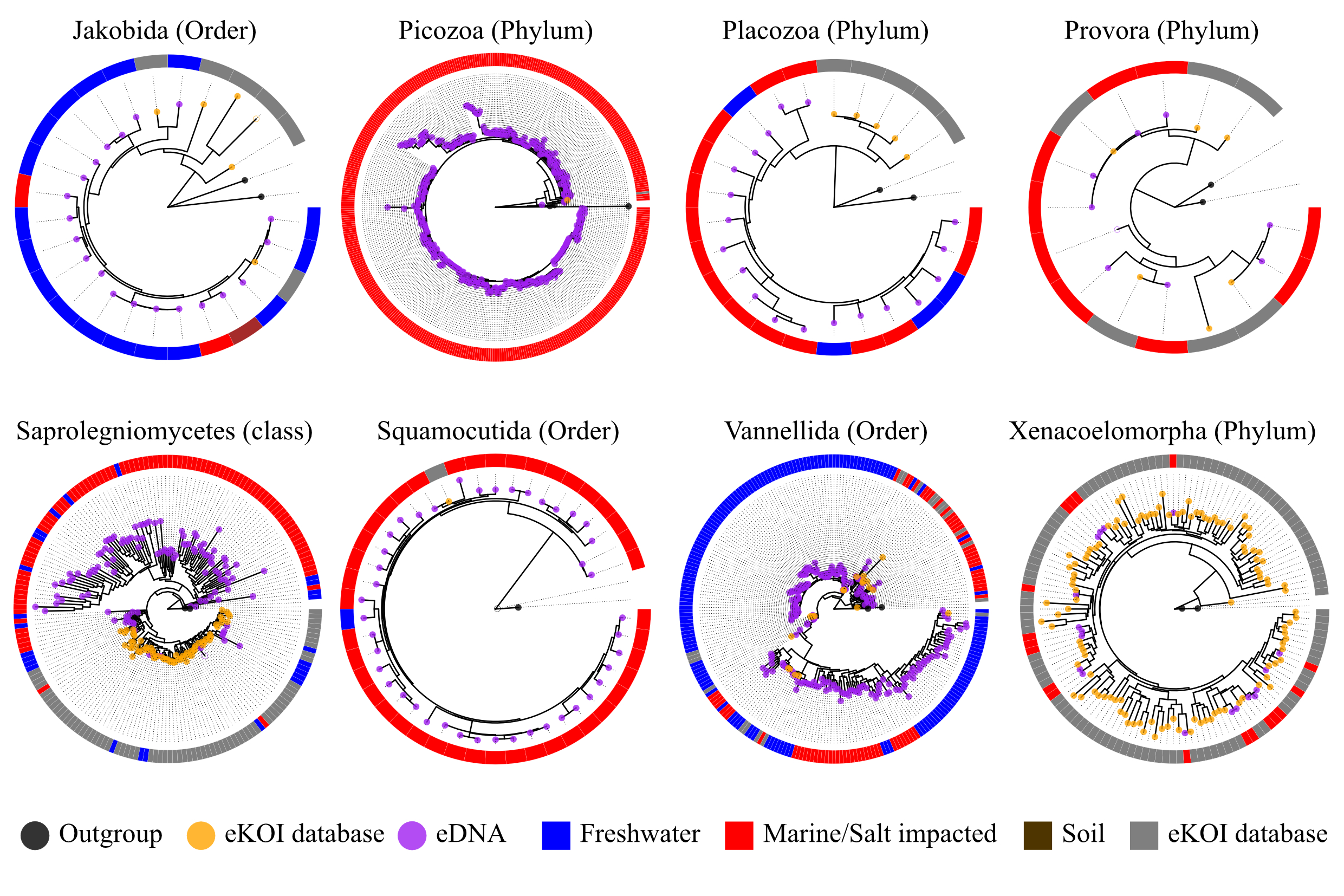

### S4

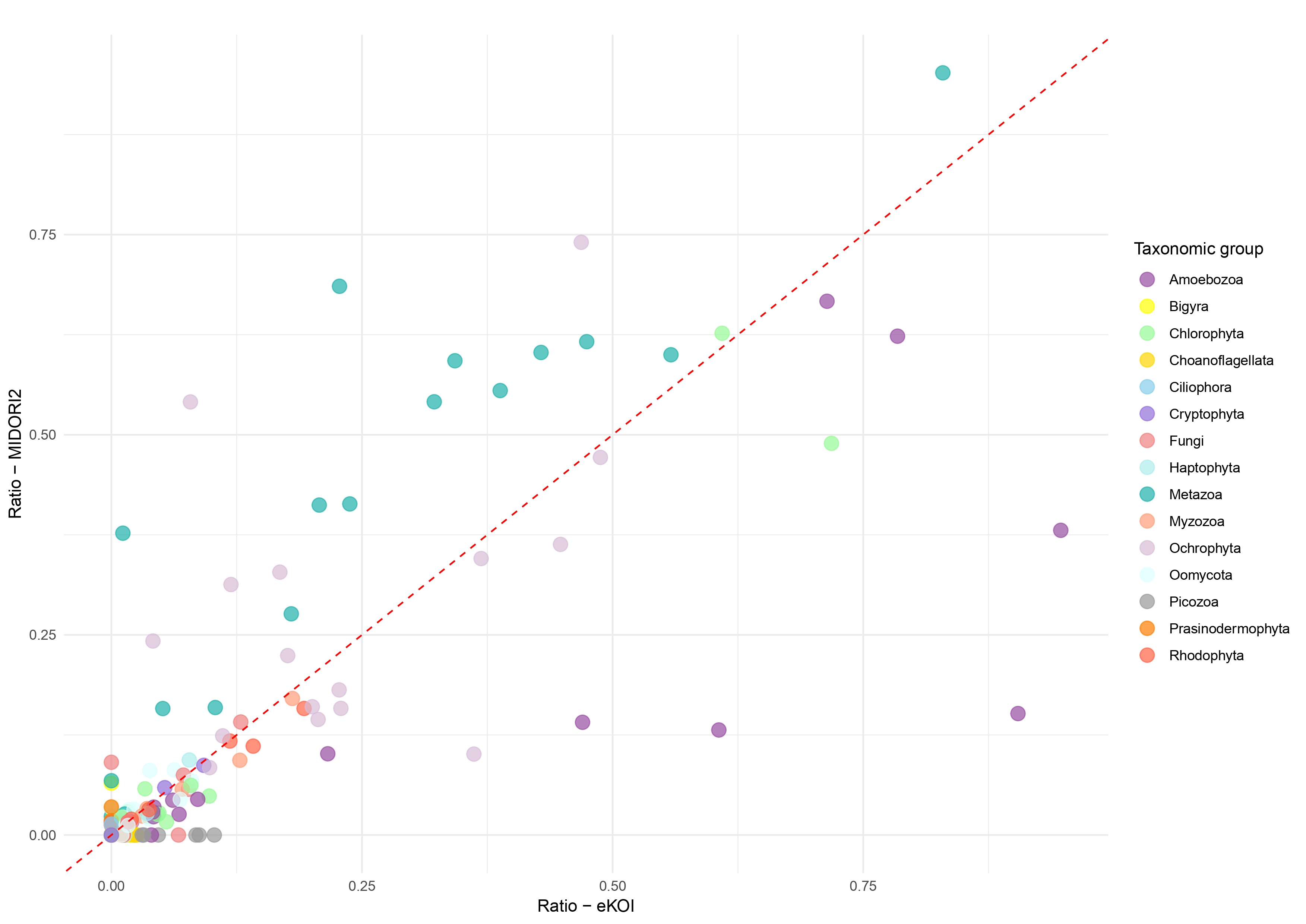
